## Supplementary Material for multiSLIDE for "multiSLIDE: a web server for exploring connected elements of biological pathways in multi-omics data"

### Table of Contents


| | |
|---|---|
| **Supplementary Figure S1** | Visualization interface of multiSLIDE |
| **Supplementary Figure S2** | Software architecture of multiSLIDE |
| **Supplementary Figure S3** | Visualization of individual omics data sets using SLIDE |
| **Supplementary Figure S4** | Pathway-based visualization of proteome and phosphoproteome data from the CPTAC Ovarian Cancer Cohort |
| **Supplementary Figure S5** | Visualization of the GO term ‘Extracellular Matrix’ in the CPTAC Ovarian Cancer Cohort |
| **Supplementary Figure S6** | Visualization of kinase-substrate relationship in the CPTAC Ovarian Cancer Cohort |
| **Supplementary Figure S7** | Distribution of correlation coefficients between kinase-site pairs in the CPTAC Ovarian Cancer Cohort |
| **Supplementary Figure S8** | Visualization of human plasma proteome and microRNA-ome associated with insulin resistance |

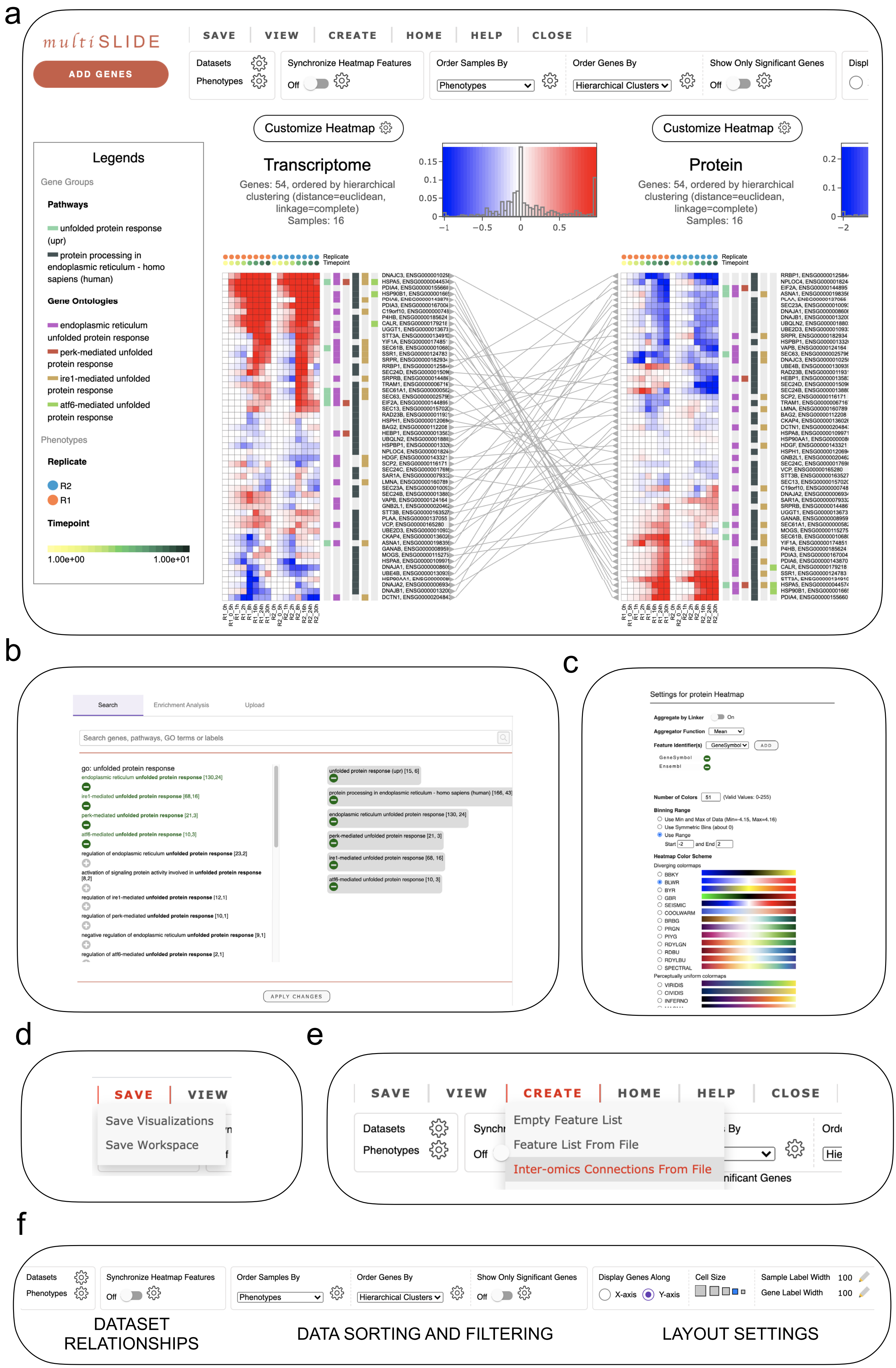

a
multiSLIDE
ADD GENES
SAVE
VIEW
CREATE
HOME
HELP
CLOSE
Datasets
Phenotypes
Synchronize Heatmap Features
Off
Order Samples By
Phenotypes
Order Genes By
Hierarchical Clusters
Show Only Significant Genes
Off
Customize Heatmap
Legends
Gene Groups
Pathways
unfolded protein response (upr)
protein processing in endoplasmic reticulum - homo sapiens (human)
Gene Ontologies
endoplasmic reticulum unfolded protein response
perk-mediated unfolded protein response
ire1-mediated unfolded protein response
atf6-mediated unfolded protein response
Phenotypes
Replicate
R2
R1
Timepoint
1.00e+00
1.00e+01
Transcriptome
Genes: 54, ordered by hierarchical clustering (distance=euclidean, linkage=complete)
Samples: 16
Protein
Genes: 54, ordered by hierarchical clustering (distance=euclidean, linkage=complete)
Samples: 16
Replicate
Timepoint
b
Search
Enrichment Analysis
Upload
Search genes, pathways, GO terms or labels
go: unfolded protein response
endoplasmic reticulum unfolded protein response [130,24]
ire1-mediated unfolded protein response [68,16]
perk-mediated unfolded protein response [21,3]
atf6-mediated unfolded protein response [10,3]
regulation of endoplasmic reticulum unfolded protein response [23,2]
activation of signaling protein activity involved in unfolded protein response [8,2]
regulation of ire1-mediated unfolded protein response [12,1]
regulation of perk-mediated unfolded protein response [10,1]
negative regulation of endoplasmic reticulum unfolded protein response [9,1]
regulation of atf6-mediated unfolded protein response [2,1]
unfolded protein response (upr) [15, 6]
protein processing in endoplasmic reticulum - homo sapiens (human) [166, 43]
endoplasmic reticulum unfolded protein response [130, 24]
perk-mediated unfolded protein response [21, 3]
ire1-mediated unfolded protein response [68, 16]
atf6-mediated unfolded protein response [10, 3]
APPLY CHANGES
c
Settings for protein Heatmap
Aggregate by Linker
On
Aggregator Function
Mean
Feature Identifier(s)
GeneSymbol
ADD
GeneSymbol
Ensembl
Number of Colors
51
(Valid Values: 0-255)
Binning Range
Use Min and Max of Data (Min=-4.15, Max=4.16)
Use Symmetric Bins (about 0)
Use Range
Start
-2
and End
2
Heatmap Color Scheme
Diverging colormaps
BBKY
BLWR
BYR
GBR
SEISMIC
COOLWARM
BRBG
PRGN
PIYG
RDYLGN
RDBU
RDYLBU
SPECTRAL
Perceptually uniform colormaps
VIRIDIS
CIVIDIS
INFERNO
d
SAVE
VIEW
Save Visualizations
Save Workspace
e
SAVE
VIEW
CREATE
HOME
HELP
CLOSE
Datasets
Phenotypes
Empty Feature List
Feature List From File
Inter-omics Connections From File
f
Datasets
Phenotypes
Synchronize Heatmap Features
Off
Order Samples By
Phenotypes
Order Genes By
Hierarchical Clusters
Show Only Significant Genes
Off
Display Genes Along
X-axis
Y-axis
Cell Size
Sample Label Width
100
Gene Label Width
100
DATASET RELATIONSHIPS
DATA SORTING AND FILTERING
LAYOUT SETTINGS

**Supplementary Figure S1**: **Visualization interface of multiSLIDE.** A. The main visualization home for multiSLIDE. On the top-left corner is the “Add Genes” button to search and add pathways and GO terms for visualization. Clicking this button opens the search panel shown in b. The three tabs in this panel, Search, Enrichment Analysis, and Upload, are respectively, for keyword-based pathway search, pathway enrichment analysis, and for uploading externally curated pathways. The menu panel at the top has options to save visualizations and analysis workspace, as well as for creating custom networks to visualize inter-omics relationships, as shown in d and e, respectively. The global settings panel, just below the menu panel, provides all the data exploration functionalities, such as joint (synchronous) or independent (asynchronous) clustering, data sorting and filtering functionalities, as shown in f. Each heatmap can be individually customized by clicking the ‘Customize Heatmap’ option beside the heatmaps, which opens the panel shown in c.

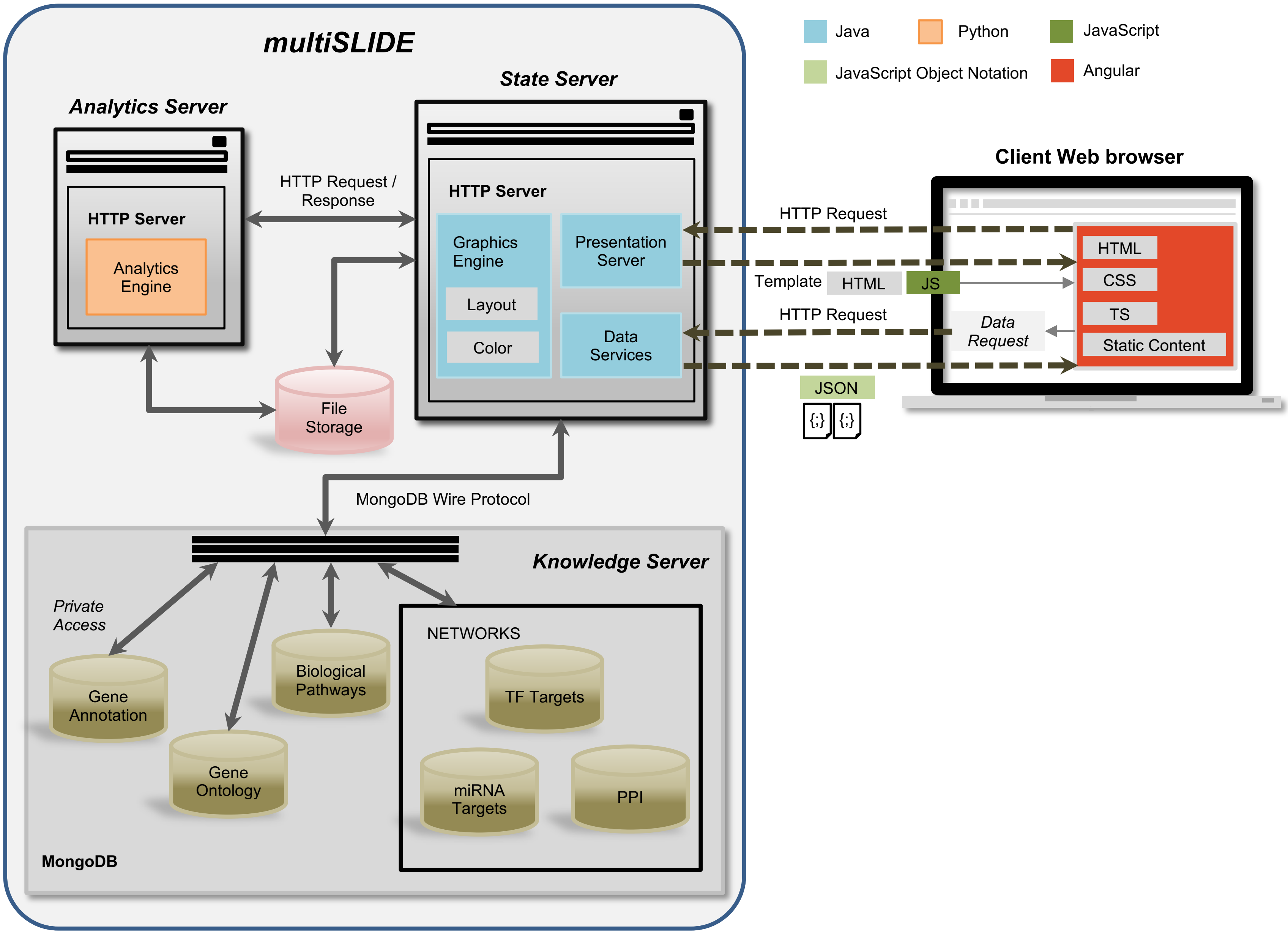
multiSLIDE
Java
Python
JavaScript
JavaScript Object Notation
Angular
State Server
Analytics Server
HTTP Server
Analytics Engine
HTTP Request / Response
HTTP Server
Graphics Engine
Presentation Server
Layout
Color
Data Services
File Storage
Client Web browser
HTTP Request
HTML
CSS
Template
HTML
JS
TS
HTTP Request
Data Request
Static Content
JSON
{;}
{;}
MongoDB Wire Protocol
Knowledge Server
Private Access
NETWORKS
Gene Annotation
Biological Pathways
TF Targets
Gene Ontology
miRNA Targets
PPI
MongoDB

**Supplementary Figure S2: Software architecture of multiSLIDE.** The figure presents a schematic of multiSLIDE's software architecture. multiSLIDE is built on a distributed architecture with the server comprising of three components: a state server, an analytics server, and a knowledge server. The client can be any modern web browser. The state and analytics servers are HTTP servers, with the state server maintaining client state information and holding uploaded data, whereas the analytics server is the main computation engine. The knowledge server implemented using MongoDB, manages the physical storage of curated gene annotation, regulatory networks, biological pathways, and Gene Ontologies (GO). The analytics and knowledge servers are stateless. The client interacts only with the state server.

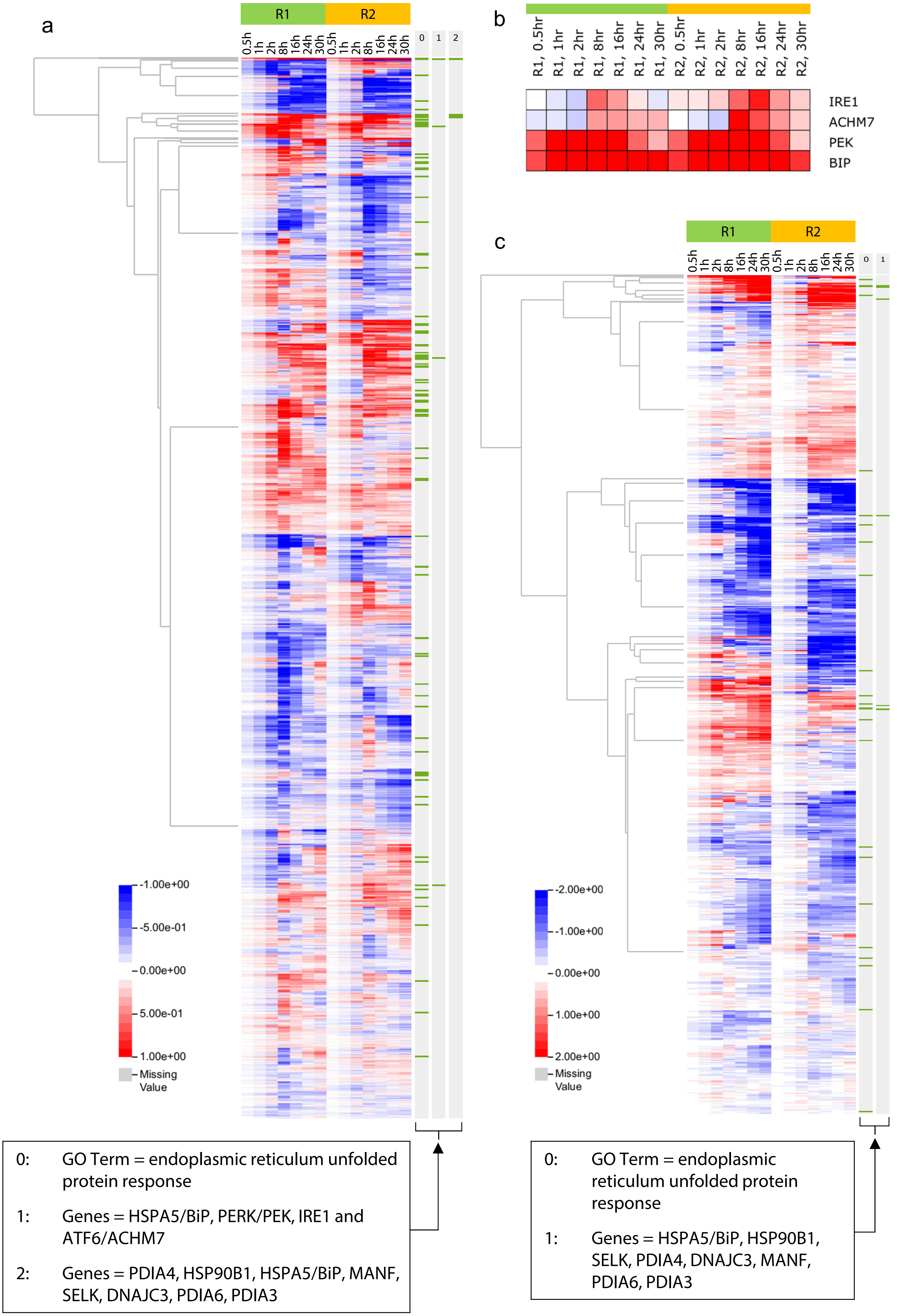

a
R1
R2
0.5h
1h
2h
8h
16h
24h
30h
0
1
2
-1.00e+00
-5.00e-01
0.00e+00
5.00e-01
1.00e+00
Missing Value
b
R1, 0.5hr
R1, 1hr
R1, 2hr
R1, 8hr
R1, 16hr
R1, 24hr
R1, 30hr
R2, 0.5hr
R2, 1hr
R2, 2hr
R2, 8hr
R2, 16hr
R2, 24hr
R2, 30hr
IRE1
ACHM7
PEK
BIP
c
-2.00e+00
-1.00e+00
0.00e+00
1.00e+00
2.00e+00
0: GO Term = endoplasmic reticulum unfolded protein response
1: Genes = HSPA5/BiP, PERK/PEK, IRE1 and ATF6/ACHM7
2: Genes = PDIA4, HSP90B1, HSPA5/BiP, MANF, SELK, DNAJC3, PDIA6, PDIA3
0: GO Term = endoplasmic reticulum unfolded protein response
1: Genes = HSPA5/BiP, HSP90B1, SELK, PDIA4, DNAJC3, MANF, PDIA6, PDIA3

**Supplementary Figure S3: Visualization of individual omics data sets using SLIDE**. a. The whole transcriptome data across the two replicates (N=16704), visualized using SLIDE's global heatmap view. The first search result highlights genes belonging to the GO term: endoplasmic reticulum (ER) unfolded protein response. The second search result highlights the four major sensor proteins in endoplasmic reticulum stress: HSPA5/BiP, PERK/PEK, IRE1, and ATF6/ACHM7. The third search result shows some known ER stress related and resident proteins. b. Visualizes the transcriptome level expression for the four sensor proteins separately. c. Visualizes the protein expression data after statistical filtering across the two replicates (N=1237) in SLIDE. The first search result highlights proteins belonging to the GO term: endoplasmic reticulum unfolded protein response. Both -omics data were baseline transformed by subtracting their respective 0 h time points (in log scale, base 2). The heatmaps show three phases of the stress response early (< 2 h), intermediate (2 - 8 h), and late (> 8 h). The whole mRNA regulation suggests a spike-like pattern in the transition from the early phase to the intermediate phase of the stress response, peaking in the intermediate phase before returning to original abundance levels in the late phase. The protein level visualization shows a delayed response that persisted till the late stage (30 h).

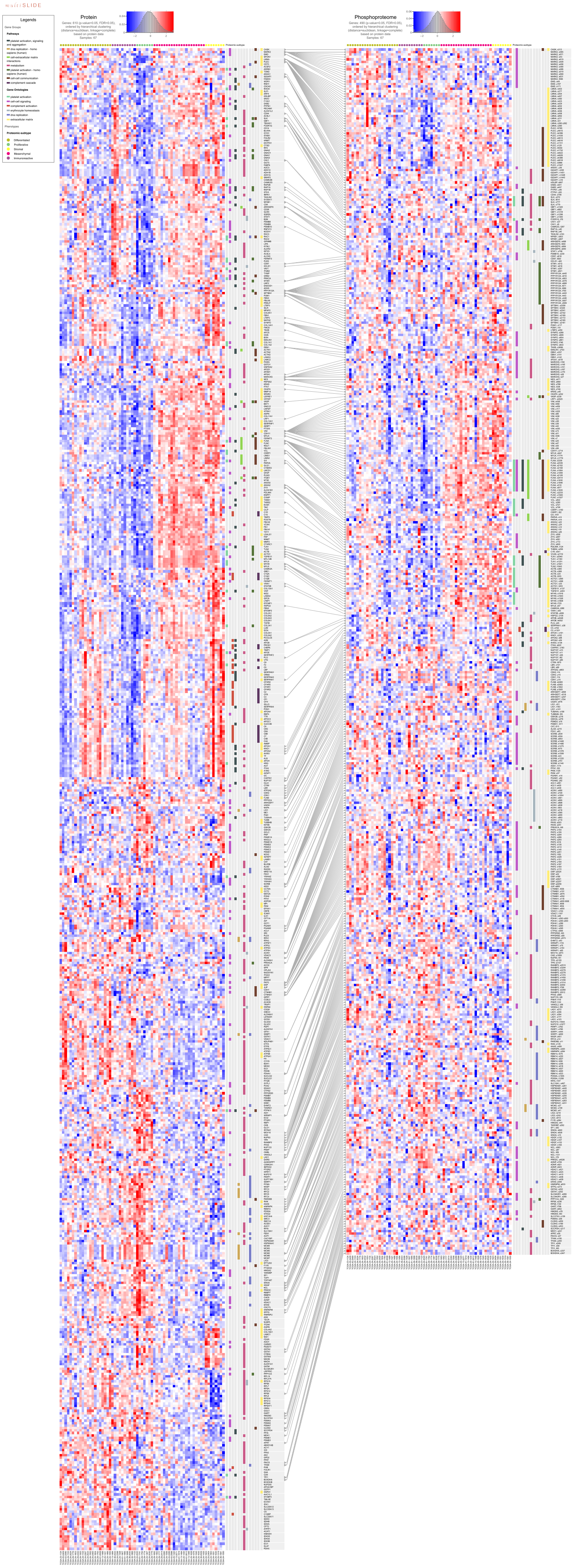

multiSLIDE
Legends
Gene Groups
Pathways
platelet activation, signaling and aggregation
dna replication - homo sapiens (human)
cell-extracellular matrix interactions
metabolism
platelet activation - homo sapiens (human)
cell-cell communication
complement cascade
Gene Ontologies
platelet activation
cell-cell signaling
complement activation
erythrocyte homeostasis
dna replication
extracellular matrix
Phenotypes
Proteomic-subtype
Differentiated
Proliferative
Stromal
Mesenchymal
Immunoreactive
Protein
Genes: 610 (p-value=0.05, FDR=0.05), ordered by hierarchical clustering (distance=euclidean, linkage=complete) based on protein data
Samples: 67
Phosphoproteome
Genes: 490 (p-value=0.05, FDR=0.05), ordered by hierarchical clustering (distance=euclidean, linkage=complete) based on protein data
Samples: 67
Proteomic-subtype

**Supplementary Figure S4: Pathway-based visualization of proteome and phosphoproteome data from the CPTAC Ovarian Cancer Cohort.** Here, the proteome-based molecular subtypes: Differentiated, Immunoreactive, Proliferative, Mesenchymal, and Stromal, are used to group samples in both the proteome and phosphoproteome data. We performed keyword-based pathway or GO term searches following the pathway enrichment analysis by Zhang et al. The legend panel shows the list of pathways and GO terms included in the visualization. Differential expression analysis using multiSLIDE's filtering feature identified 610 proteins and 490 phosphosites, using one-way ANOVA (p-value <= 0.05) for molecular subtypes and 5% FDR cutoff, as indicated beside the histograms. The heatmaps are synchronized, and hierarchical clustering of the protein data is used to determine the order. The visualization shows that there are similar expression patterns across the two omics levels, with phosphosites predominantly upregulated in the Mesenchymal and Stromal subtypes.

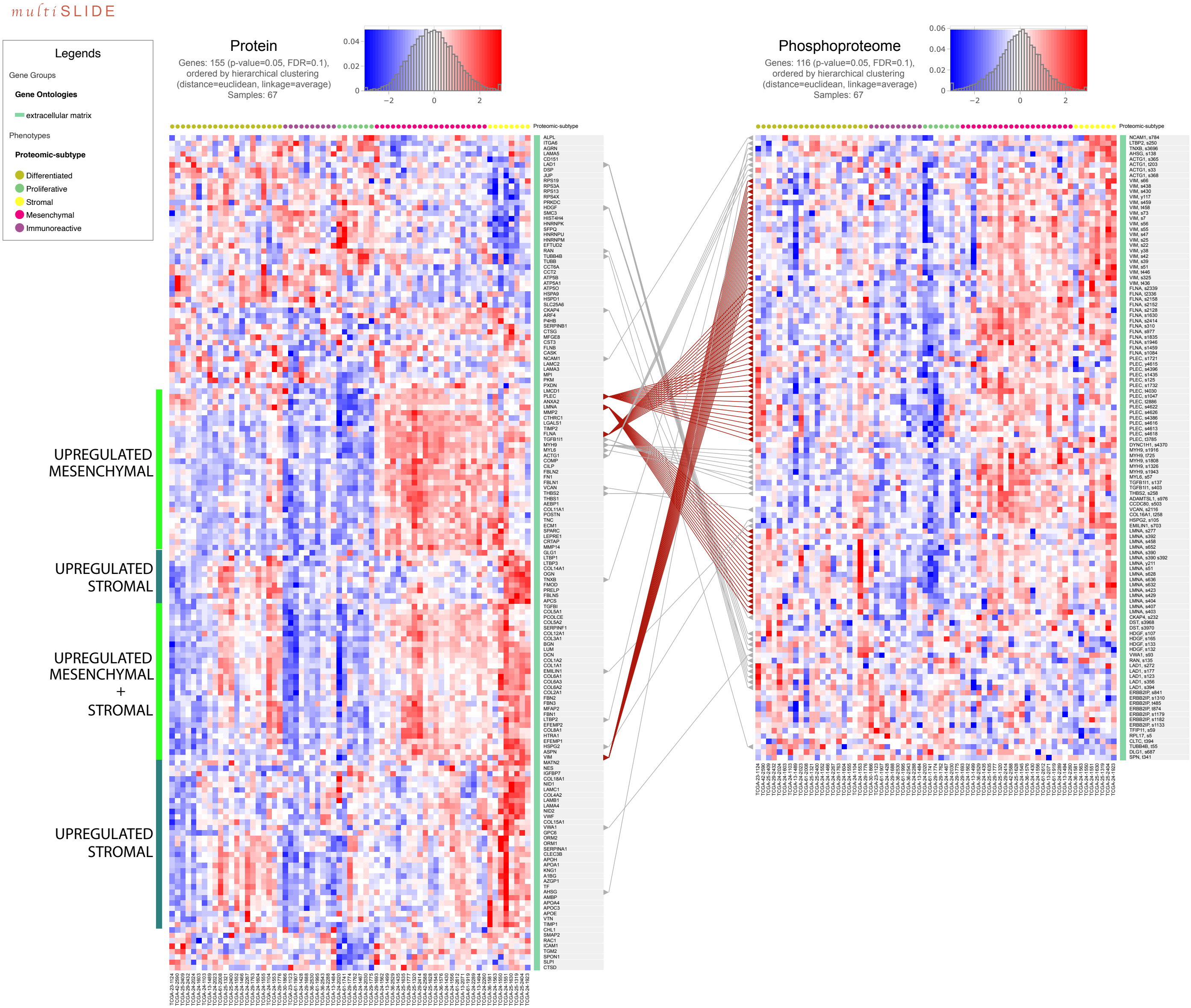

multiSLIDE
Legends
Gene Groups
Gene Ontologies
extracellular matrix
Phenotypes
Proteomic-subtype
Differentiated
Proliferative
Stromal
Mesenchymal
Immunoreactive
Protein
Genes: 155 (p-value=0.05, FDR=0.1),
ordered by hierarchical clustering
(distance=euclidean, linkage=average)
Samples: 67
Phosphoproteome
Genes: 116 (p-value=0.05, FDR=0.1),
ordered by hierarchical clustering
(distance=euclidean, linkage=average)
Samples: 67
Proteomic-subtype
UPREGULATED MESENCHYMAL
UPREGULATED STROMAL
UPREGULATED MESENCHYMAL + STROMAL
UPREGULATED STROMAL

**Supplementary Figure S5: Visualization of the GO term ‘Extracellular Matrix’ in the CPTAC Ovarian Cancer Cohort**. Genes belonging to the GO term “Extracellular Matrix” are visualized here. Differential expression analysis using multiSLIDE's filtering feature identified 155 proteins and 116 phosphosites. The filtering used one-way ANOVA (p-value <= 0.05) based on the molecular subtypes and 10% FDR cutoff. In contrast to **Supplementary Figure S4**, here, proteins and phosphosites are clustered independently. The clusters in the protein data are highlighted using green bars. The visualization shows that proteins such as Plectin (PLEC), Lamin A/C (LMNA), Filamin A (FLNA), and Vimentin (VIM) have the highest number of sites that are differentially phosphorylated, with consistent patterns with the differences in the protein abundance.

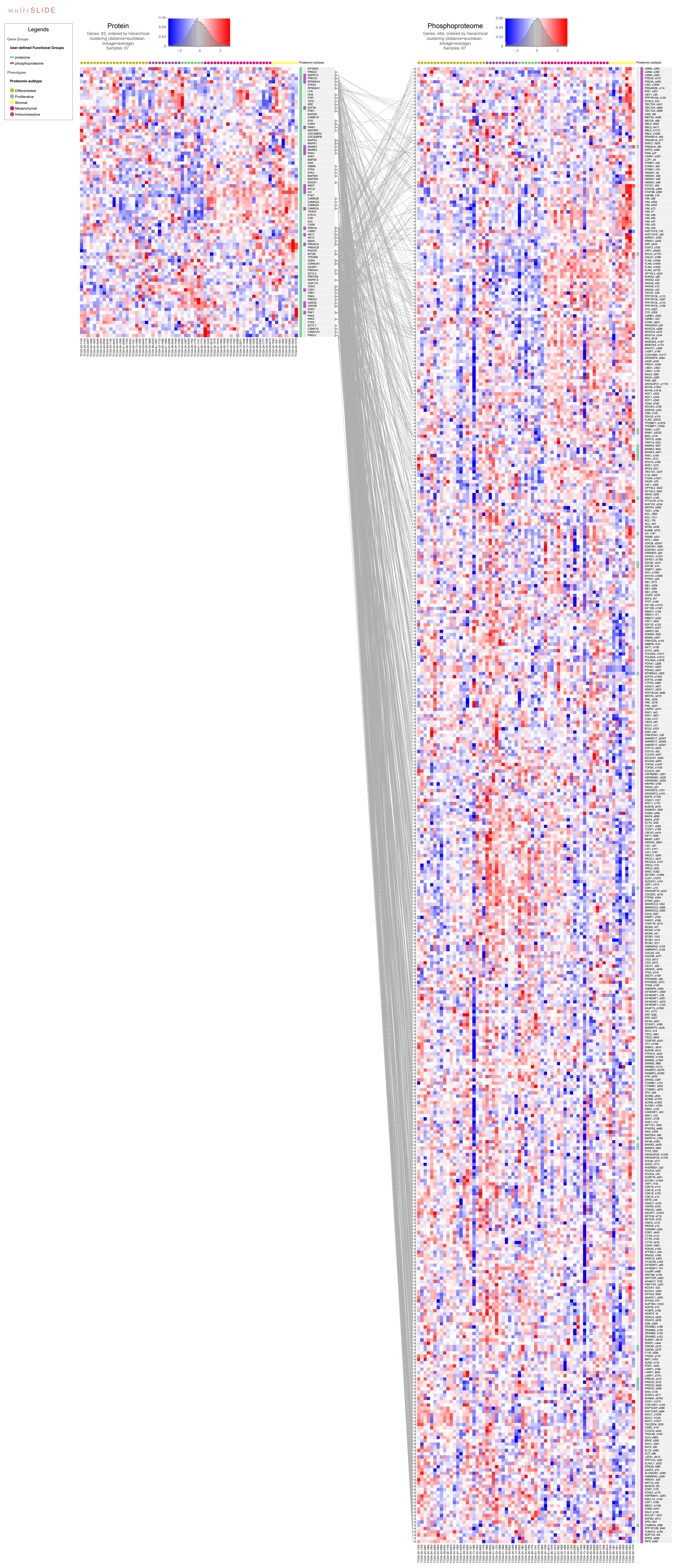
multiSLIDE
Legends
Gene Groups
User-defined Functional Groups
proteome
phosphoproteome
Phenotypes
Proteomic-subtype
Differentiated
Proliferative
Stromal
Mesenchymal
Immunoreactive
Protein
Genes: 83, ordered by hierarchical clustering (distance=euclidean, linkage=average)
Samples: 67
Phosphoproteome
Genes: 454, ordered by hierarchical clustering (distance=euclidean, linkage=average)
Samples: 67
Proteomic-subtype

**Supplementary Figure S6: Visualization of kinase-substrate relationship in the CPTAC Ovarian Cancer Cohort**. The figure visualizes kinases in the proteome data and their corresponding substrates in the phosphoproteome data. We curated kinase-substrate interactions from PhosphoSitePlus (1), PhosphoNetworks (2), and a predictive network inference approach (3) to build a kinase-substrate map. All kinase-substrate pairs present in both the kinase-substrate map and in the datasets are visualized here. Figure 3 shows a subset of these kinases-substrate pairs, where the proliferative subtype is upregulated.

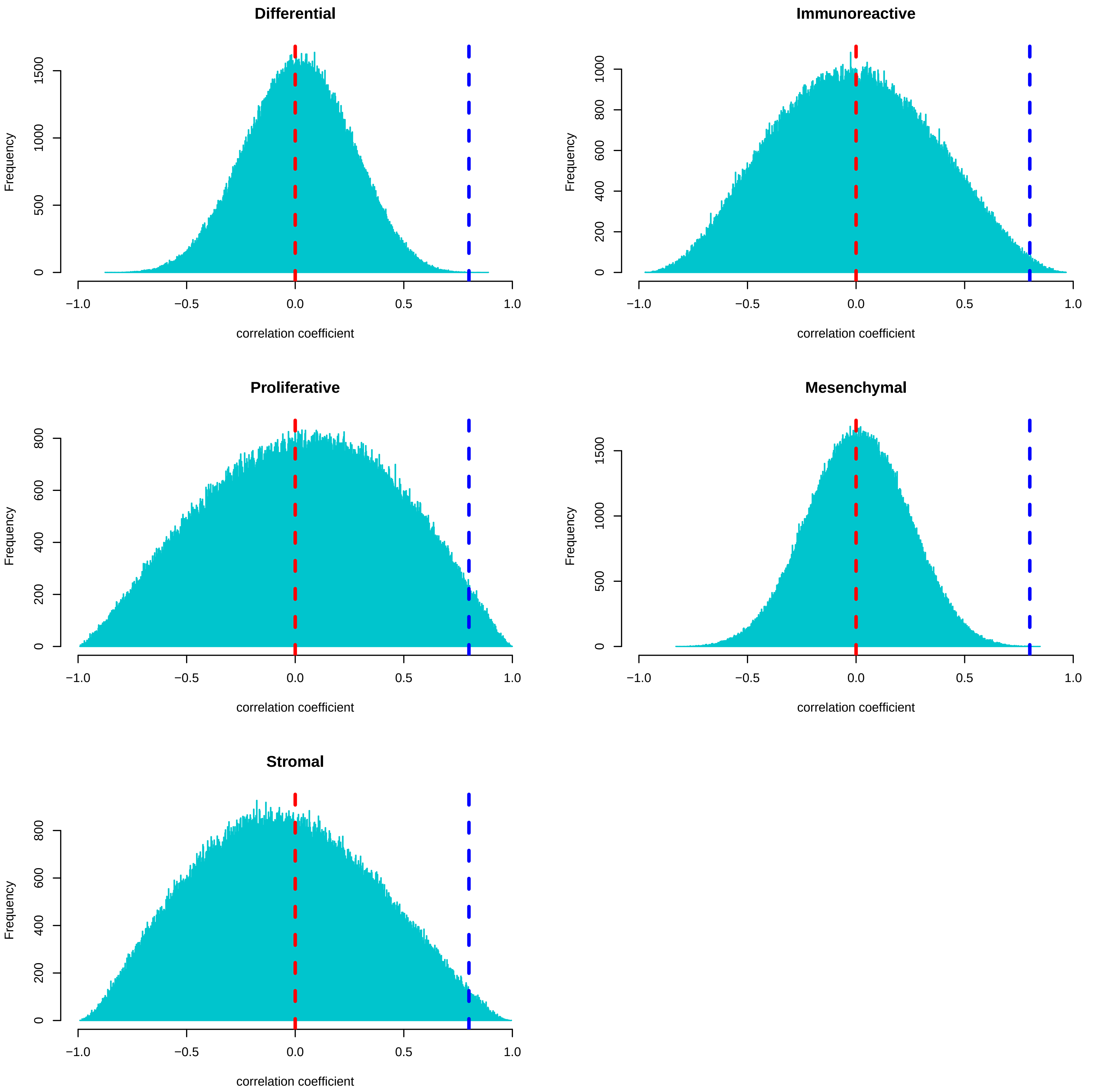
Differential
Immunoreactive
Proliferative
Mesenchymal
Stromal
Frequency
correlation coefficient
−1.0
−0.5
0.0
0.5
1.0
0
500
1000
1500
200
400
600
800

**Supplementary Figure S7: Distribution of correlation coefficients between kinase-site pairs in the CPTAC Ovarian Cancer Cohort**. The five panels show the distribution of the Pearson correlation coefficients between the subtype-specific expressions of kinase-site pairs for the five molecular subtypes in the CPTAC Ovarian Cancer Cohort. The correlation coefficients are calculated between the 83 kinases visualized in **Supplementary Figure S6** and all phosphosites. The blue vertical line indicates a correlation value of 0.8. For the Immunoreactive, Proliferative, and Stromal subtypes, a large number of positively correlated kinase-site pairs are present. We also see in **Supplementary Figure S6** that several kinases are upregulated in the Proliferative subtype. Therefore, in Figure 3, we focus on kinase-site pairs that are highly correlated and show upregulation in the Proliferative subtype.

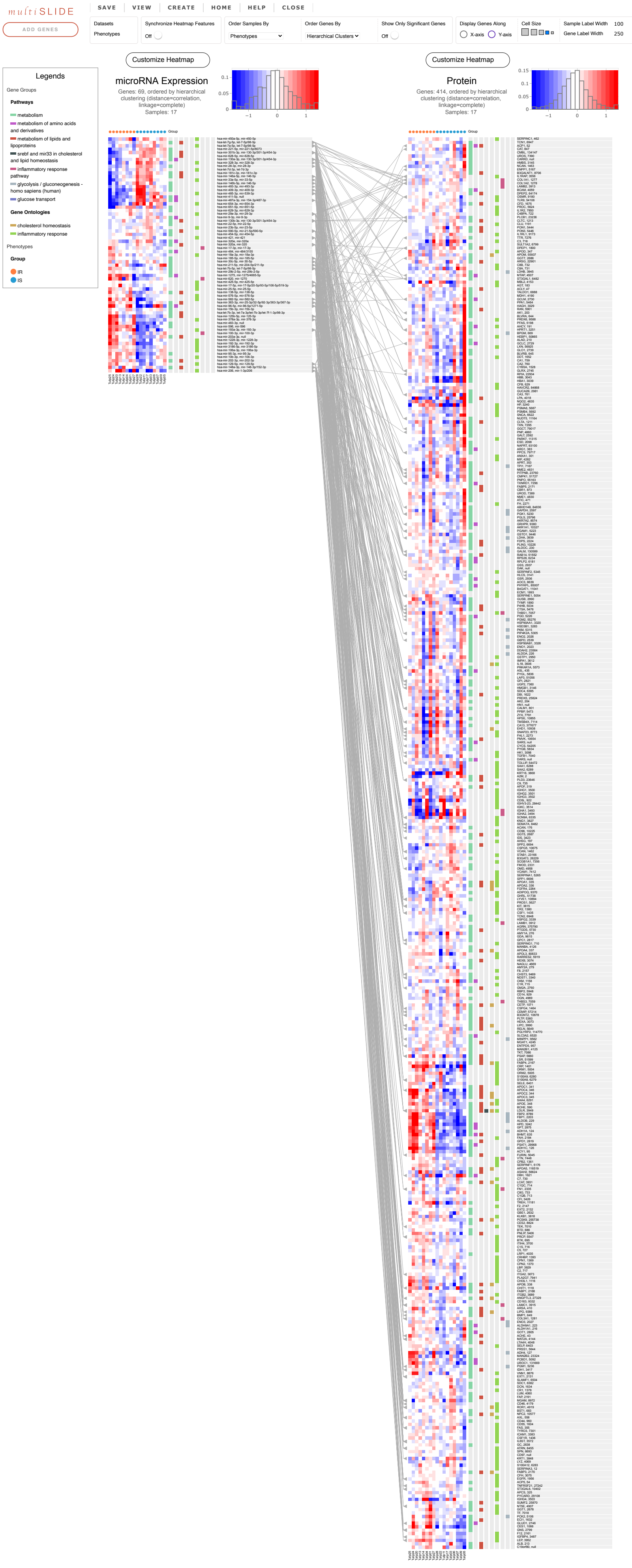
multiSLIDE
ADD GENES
SAVE
VIEW
CREATE
HOME
HELP
CLOSE
Datasets
Phenotypes
Synchronize Heatmap Features
Off
Order Samples By
Phenotypes
Order Genes By
Hierarchical Clusters
Show Only Significant Genes
Off
Display Genes Along
X-axis
Y-axis
Cell Size
Sample Label Width
100
Gene Label Width
250
Customize Heatmap
microRNA Expression
Genes: 69, ordered by hierarchical clustering (distance=correlation, linkage=complete)
Samples: 17
Customize Heatmap
Protein
Genes: 414, ordered by hierarchical clustering (distance=correlation, linkage=complete)
Samples: 17
Legends
Gene Groups
Pathways
metabolism
metabolism of amino acids and derivatives
metabolism of lipids and lipoproteins
srebf and mir33 in cholesterol and lipid homeostasis
inflammatory response pathway
glycolysis / gluconeogenesis - homo sapiens (human)
glucose transport
Gene Ontologies
cholesterol homeostasis
inflammatory response
Phenotypes
Group
IR
IS
Group

**Supplementary Figure S8: Visualization of human plasma proteome and microRNA-ome associated with insulin resistance.** The figure shows pathways selected by searching for keywords: metabolism, inflammatory response, glucose transport, and lipid homeostasis. Here, proteins and miRNA are independently clustered using correlation distance (1 minus Pearson correlation) and complete linkage function. The relationships between miRNA family names and their target proteins are extracted from TargetScanMap and uploaded into multiSLIDE using the network upload feature. A subset of these proteins and miRNA, identified through differential expression analysis, are visualized in **Figure 4**.
